# EFFECT OF DRY SANITIZATION ON DRY SURFACE BIOFILM OF *Cronobacter sakazakii*

**DOI:** 10.64898/2026.08.05.742776

**Authors:** Raul Fernando Pereira, Vinícius S. A. Vaz, Rafael Pimentel Maia, Jean-Yves Maillard, Maristela da Silva do Nascimento

## Abstract

*Cronobacter sakazakii* is an opportunistic foodborne pathogen that affects neonates, and has ability to produce dry surface biofilm (DSB). However, little is known about its resistance to dry sanitizers. This study aimed to evaluate the efficiency of dry sanitizers compared with sodium hypochlorite (SH) on DSB of *C. sakazakii*. DSB were formed on stainless steel or polypropylene coupons (10 cm²) using two cycles of hydrated/dry phases at 25 °C: T1 (48/48 h) or T2 (24/120 h). At DSB endpoint, the coupons were sanitized with 70% ethanol, a commercial product based on isopropyl alcohol (25%) and quaternary ammonium (0.015%), hot air (90±2 °C), UV-C light (254 nm), gaseous ozone (45±2 mg/L), or SH (200 mg/L, pH 6.5). All dry sanitizers showed limited antimicrobial activity (p > 0.05), with reductions ≤ 0.8 log CFU/cm² over 30 min exposure. In contrast, SH was effective against DSB regardless of the DSB formation protocol, with reductions ≥2 and >5 log CFU/cm² after 10 and 30 min, respectively. Interestingly, DSB formed with shorter hydrated phase and longer dry phase (T2) had greater sensitivity to SH, but it was not noted to dry sanitizers. CLSM images suggested the presence of VBNC cells, particularly after SH treatment. In conclusion, our results emphasize the importance of implementing stringent hygiene measures to control *C. sakazakii* DSB in the low moisture food industry, and indicate that SH is an effective sanitation strategy when the drying line is promptly addressed.

## 1. Introduction

*Cronobacter sakazakii* is characterized as an opportunistic pathogen that affects vulnerable populations such as neonates and immunocompromised elderly (Farmer et al., 1980; Mousavi et al., 2023). Donaghy et al, (2025) classify this pathogen as of higher risk for children under 6 months, especially infants less than 60 days old. In recent years, two episodes of infection in newborns linked to powdered infant formula (PIF) contaminated with *C. sakazakii* have been reported in the US, one isolated case in 2021 and an outbreak in 2022, with four cases and two deaths (CDC, 2025; Haston et al., 2023). Furthermore, around the world between 2022 and 2024, there were seven cases of recalls/withdrawals/outbreaks involving PIF contaminated with *C. sakazakii* (Donaghy et al., 2025). The worldwide prevalence of *C. sakazakii* in powdered milk and flours is approximately 8.4%; in Brazil, the prevalence in PIF (powdered milk) was 23.33% (Ekundayo & Ijabadeniyi, 2024). Furthermore, the pathogen has already been isolated from chestnuts, rice, corn, wheat, and other cereals and flours (Samadpour et al., 2024, Lou et al., 2019). *C. sakazakii* has ability to form biofilms on different surfaces, such as polyvinyl chloride (PVC), polypropylene (PP) and stainless steel (SS) (Gemba et al., 2025). According to Umeda et al. (2017), several strains of *C. sakazakii,* isolated from food and clinical samples in Brazil, are able to form biofilms on polystyrene plates. *C. sakazakii* is highly resistant to desiccation, and was recovered from PIF samples after 90 days (Hennekinne et al., 2018). This ability may be related to metabolic and physiological changes in biofilm cells (Lou et al., 2024). The physiological changes occur initially by the accumulation of substances that provide protection against external osmotic pressure, such as potassium and glutamate. In a second stage, the synthesis of solutes occurs, such as saccharides, free amino acids, small peptides, sulfate esters; there is a five-fold increase in the trehalose concentration (Wang et al., 2024). This resistance not only facilitates the survival of *C. sakazakii*, but also hinders its eradication from low moisture food (LMF) processing facilities (Breeuwer et al., 2003).

Stressful conditions such as desiccation, heat treatment, lack of nutrients or presence of antimicrobials can trigger biofilm formation as an adaptive response of the microorganisms (Venkatesan & Sikkander, 2023). Biofilm adhesion is influenced by the type of surface and its characteristics, such as topography, texture, and the presence of cracks or crevices, as well as moisture content (Faille et al., 2018). Among biofilms, there is one type that requires greater attention from the LMF industry, the dry surface biofilm (DSB). This type of biofilm was first reported in the medical area by Vickery et al. (2012), and more recently it has been studied in foodborne pathogens (Duggan et al., 2024; Chaggar et al., 2024; Lin et al., 2024). For DSB formation, alternating hydrated and dry phases is necessary. Bacteria need a stage with high moisture to enable its growth, followed by a dry step that activates its responses to desiccation stress. Studies have suggested that these biofilms may be resistant to sanitization and that some pathogens may reduce their metabolism and survive for months in this condition (Maillard & Centeleghe, 2023; Chaggar et al., 2024). In general, DSB EPS has a lot of protein, a moderate amount of carbohydrate, and little DNA (Ledwoch et al., 2019).

To prevent biofilm formation, wet sanitization is routinely employed in the food industry. Among the main sanitizers used is sodium hypochlorite (SH), whose active principle is the hypochlorous acid (HOCl) (Gurtler et al., 2014). The concentration of this acid in its undissociated form is governed by pH, and it is predominant at values < 7.0. HOCl is a type of reactive oxygen species (ROS), that is, a potent oxidizing agent capable of causing loss of cell membrane transport, disruption of the electron transport chain, dissipation of adenylate energy reserves, and suppression of DNA synthesis (Falk et al., 2020). SH at 250 mg/L caused reductions of around 3 log CFU/cm² in *C. sakazakii* hydrated biofilms formed on PP and SS (Bayoumi et al., 2012). Regarding its performance against *Candida auris* and *Staphylococcus aureus* DSB, reductions of 2 to >7 log CFU/cm² were obtained using concentrations of 500 mg/L and 10,000-20,000 mg/L, respectively (Ledwoch & Maillard, 2018; Almatroudi et al., 2016). However, wet cleaning is not recommended for LMF industry. Water entry into the factory environment should be restricted, and dry sanitizing alternatives should be employed to prevent microbial growth (Codex Alimentarius, CXC 75 2018). Among the dry sanitizers are alcohol-based, hot air, UV-C and ozone. While alcohol-based compounds act by denaturing and/or coagulating proteins and affecting cell membrane and enzymes (Kampf & Arbogast, 2020), hot air chemically oxidizes and dehydrates the components of bacterial cells (Alonso et al., 2022). UV-C radiation absorbed by the cell generates photoproducts that damage DNA (Blatchley III & Coohill, 2020). Ozone is also a potent oxidizing agent, which can generate other short-lived oxidizing agents, such as hydroxyl radicals, resulting in degradation of genetic material (Weavers et al., 2020).

Due to the prevalence of *C. sakazakii* in powdered infant formula and its high pathogenicity in neonates, the FDA created a strategy to assist in the control and prevention of this pathogen (FDA, 2023). For that, it is necessary to broaden scientific knowledge and understanding on *C. sakazakii*, especially in low moisture environment. However, to date there are no studies evaluating the resistance of *C. sakazakii* DSB to chemical or physical sanitizers. Therefore, this study is a pioneer in investigating the efficiency of dry sanitizers on DSB formed by *C. sakazakii* on polypropylene (PP) and stainless steel (SS).

## 2. Material and methods

### 2.1. Origin of isolates and preparation of inoculum

A pool of five strains of *C. sakazakii* was used in this study. They were previously isolated from low moisture products (P4499 - isolated from pre-cooked milk flour; P4787 - isolated from pre-cooked rice-based cereal; P4791 - ground ginger; P4795 - oat and rice porridge; and P4798 - breadcrumbs) (Brandão et al., 2017). In addition, all strains had previously demonstrated the ability to form hydrated biofilms (Umeda et al., 2017). The strains were stored at -80°C in an ultralow freezer (NuAire, NU-6518G, MN, USA) and recovered in brain and heart infusion (BHI, Difco, MD, USA) broth with incubation for 18-20 h at 37 °C. Subsequently, the strains were streaked on trypticase soy agar (TSA, Difco) and incubated for 24 h at 37 °C. To obtain the final inoculum, suspensions of each strain were prepared in saline solution (0.85%) (Synth, Brazil) to achieve turbidity equivalent to 0.5 McFarland. Then, 1 mL of each strain suspension was transferred to a sterile tube to form a stock suspension. After vortexing for 30 s at 30,000 rpm, the suspension was diluted in trypticase soy broth (TSB, Difco) to achieve a final concentration of ca. 6 log CFU/mL.

### 2.2. Surface preparation

PP and AISI 304 SS coupons measuring 2 x 5 cm were used as surface materials to form DSB. Initially, the coupons were washed with neutral detergent and rinsed with distilled water, followed by an ultrasonic bath at 40 kHz for 15 min (Ultronique, Brazil). Finally, the coupons were sterilized in an autoclave for 30 min at 121 °C and dried in an oven at 50 °C.

### 2.3. DSB formation

The study evaluated two DSB formation protocols, each consisting of two hydrated/dry cycles. Protocol T1 comprised a 48-h hydrated phase and a 48-h dry phase, and protocol T2 comprised a 24-h hydrated phase and a 120-h dry phase (adapted from Ledwoch et al., 2019). The coupons were placed in sterile Petri dishes (9 cm diameter) containing a circular polypropylene support (8.8 cm diameter) with a rectangular opening in the center (4.3 x 6.7 cm), so that only one face of the coupons remained in contact with the wet inoculum (TSB). Each plate with a support containing three coupons was inoculated with 10 mL of the inoculum, prepared as described in section 2.1. After incubation for 24 or 48 h at 25 °C (first hydrated phase), TSB was drained, and the coupons were transferred to another Petri dish and incubated at 25 °C for 48 or 120 h (first dry phase). The coupons were then transferred again to Petri dishes containing the polypropylene supports and 10 mL TSB, without the inoculum, were carefully added and plates were incubated at 25 °C for 48 or 120 h (second hydrated phase). This was followed with a second dry phase as described above. At the end of the second dry phase (DSB endpoint), DSB on coupons were ready to be tested (Figure 1).

**Figure 1.**
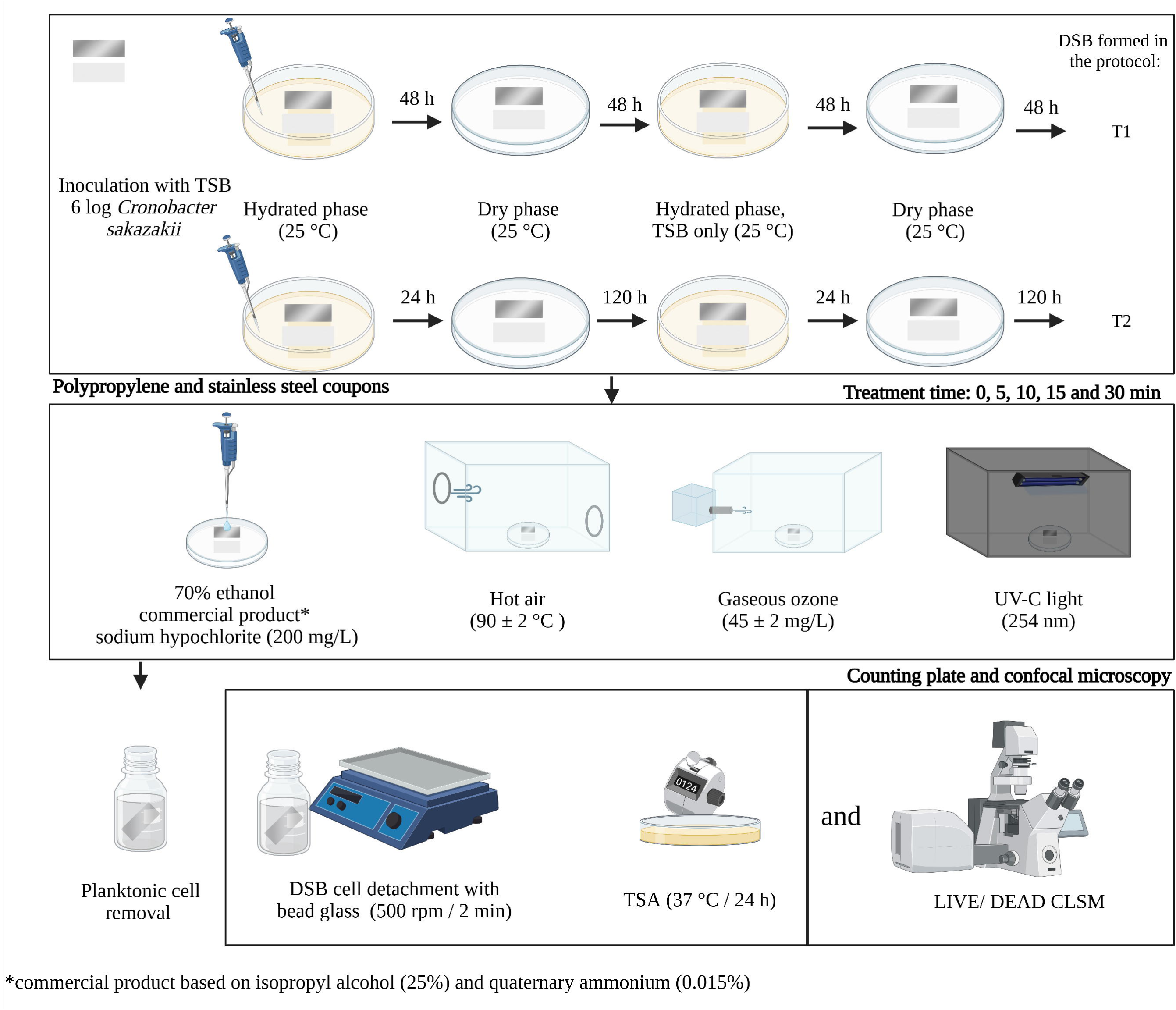
Experimental design of *Cronobacter sakazakii* dry surface biofilm (DSB) forming on polypropylene (PP) and stainless steel (SS) and sanitizing tests.

### 2.4. Sanitization

Sanitization tests were performed into a biological safety cabinet. The sanitizers evaluated were: 70% ethanol (v/v), a commercial product based on isopropyl alcohol (25%) and quaternary ammonium compound (0.015%), hot air (90 °C ± 2 °C), UV-C light (254 nm), gaseous ozone (45 ± 2 mg/L), and sodium hypochlorite (SH 200 mg/L, pH 6.5). Only the coupon face on which DSB was formed was exposed to the sanitizers for 0 (positive control), 5, 10, 15, and 30 min (adapted from Harada & Nascimento, 2021a). Coupons without DSB formation were used as negative control. Three independent tests were performed for each experiment (Figure 1).

#### 2.4.1. Alcohol-based sanitizers

The coupons were placed in a Petri dish covered with 400 μl of 70% ethanol or the commercial product. The 70% ethanol solution was prepared fresh by mixing sterile distilled water with absolute ethyl alcohol (Labsynth, Brazil). After each sanitization time, coupons of PP and SS were gently transferred to another Petri dish containing 20 mL of Letheen broth (Acumedia, MI, USA) supplemented with Tween 80 (Sigma Aldrich, Germany) and keep for 5 min to be neutralized.

#### 2.4.2. Hot Air

For hot air treatment, the coupons were placed on a glass Petri dish and transferred to a glass box (30 x 15 x 19 cm) with a small opening on the left side to which an air blower device was connected (Taiff, Brazil). The coupons were treated with a continuous flow (approximately 2.0 m/s) at 90 ± 2 °C. The temperature was monitored with a thermocouple (PT100 - Testo 112, Testo, Germany). After exposure for the appropriate contact time, coupons were removed for analysis (see 2.5 below).

#### 2.4.3. Gaseous ozone

The coupons were placed inside a hermetic acrylic box (30 x 20.5 x 20 cm), which were previously sanitized with 70% ethanol. This box was connected to an ozone generator (Ozoxi, Brazil), an ozone meter (UV-100, EcoSensors, NM, USA) and an ozone destroyer (Ozoxi, Brazil) at the other outlet. Ozone concentration used was 45 ± 2 mg/L at 20 °C. The box was placed inside an exhaust hood to prevent accidental exposure to the analyst (adapted from Nicholas et al., 2013). After exposure for the appropriate contact time, coupons were removed for analysis (see 2.5 below).

#### 2.4.4. UV-C

For UV-C treatment, the coupons were arranged in a glass petri dish and then placed in a dark box (30 x 15 x 15 cm) containing an 8W, 254nm UV-C lamp (OSRAM, Germany), 15 cm above the coupons, with a power of ca. 6.5 mW/cm², measured with a radiometer (Maestro, Gentec, Canada). After exposure for the appropriate contact time, coupons were removed for analysis (see 2.5 below).

#### 2.4.5. Sodium hypochlorite (SH)

The coupons were placed in a glass petri dish and covered with 400 μl of SH (200 mg/L - pH 6.5). The sanitizer solution was prepared and titrated (ABNT NBR 9425) immediately before use, and pH adjusted with hydrochloric acid (0.1 N; Merck, Germany) to 6.5. After each exposure time, test coupons were gently immersed in 20 mL of a 0.85% saline solution + 1% sodium thiosulfate (Labsynth, Brazil) for 5 min to neutralize the sanitizer.

### 2.5. DSB enumeration post-exposure

Prior to enumeration of survival bacteria in DSB, loosely attached planktonic cells were removed. The neutralization process described above for liquid sanitizers served to remove planktonic cells. For other treatments, planktonic cells were removed by the coupon immersion in saline solution for 5 min (adapted from Harada & Nascimento, 2021a, 2021b).

After neutralization and planktonic cell removing, the coupons were transferred to tubes containing saline solution (0.85%) and glass beads and shaken (500 rpm) for 2 min to release adhered cells (adapted from Ziech et al., 2016). Then, serial decimal dilutions were performed in peptone water (0.1% w/v) (Difco) and plated on TSA. After incubation at 37°C for 24 h, colonies were counted. The results were expressed in log CFU/cm², and the detection limit was 0.7 log CFU/cm².

### 2.6. CLSM (LIVE/DEAD)

In order to compare distribution of live and dead cells between the two DSB protocols, an untreated coupon (0 min) and a 30-min exposure coupon from each sanitizer were analyzed by CLSM. The PP and SS coupons were stained using the LIVE/DEAD Biofilm Viability Kit (Invitrogen, Eugene, OR, USA), composed of Propidium Iodide, a fluorophore that stains dead cells, and SYTO® 9, which stains living cells, both at a dilution of 1:1000. After 10 min contact, the coupons were washed with PBS solution. The images were acquired using a confocal microscope composed of an LSM 780-NLO coupled to the Axio Observer (Carl Zeiss AG, Germany) and a Zeiss EC Plan-Neofluar 20x/0.50 M27 objective lens. The images were processed using FIJI (ImageJ) software (https://imagej.net/software/fiji/) to measure the percentage of green and red cells, and the intensity of each channel.

### 2.7. Statistical analysis

To evaluate the effectiveness of sanitizers on the DSB of *C. sakazakii*, the population reduction values obtained after each sanitization time (log CFU/cm²) were analyzed by one-way ANOVA and multiple comparisons using Tukey’s test. The Shapiro-Wilk and Breusch-Pagan tests confirmed the normality and homoscedasticity of the dataset. The analyses were conducted in the R statistical environment (RStudio - R 4.5.1, 2025).

## 3. Results and Discussion

### 3.1. Efficacy of sanitizers against *C. sakazakii* DSB

This study is the first to assess the effectiveness of different dry sanitizing methods against *C. sakazakii* DSB. The FDA (2023) has identified the manufacturing environment as a potential source of PIF contamination by *C. sakazakii*. Due to the ability of this pathogen to persist in the environment for long-term (Iversen & Forsythe, 2004), it is crucial to establish effective mitigation strategies. Although wet sanitization is not recommended as the first option for PIF processing plants (Codex Alimentarius, CXC 66 2008), SH was used for comparison purposes, as it is widely used in the food industry.

To date, no official method exists for evaluating the efficacy of dry sanitizers against biofilms, nor is there a standardized protocol for assessing sanitizer performance against DSB. In the USA, the Environmental Protection Agency (EPA) provides guidance for assessing the effectiveness of wet sanitizers seeking biofilm-related claims. The agency recommends the ASTM E3161-25 method for biofilm formation and the ASTM E2871-25 method for evaluating the effectiveness of wet sanitizers against biofilms grown in the CDC biofilm reactor using the single tube method (ASTM, 2025a, 2025b). The EPA also establishes a performance criterion of at least a 6-log CFU reduction per coupon within 10 min exposure (EPA, 2025). In Europe, the UNE-EN 13697 (2024) standard focuses on assessing the efficacy of wet sanitizers on dried cells on surfaces and sets a performance criterion of ≥ 4-log CFU reduction. Ledwoch et al. (2019) adapted the ASTM E2967-15 method to evaluate the combined effect of chemical sanitizers and mechanical action—using a wiperator—on DSB formed on stainless steel coupons by the sedimentation method. However, it is important to emphasize that no official methodology for *in vitro* DSB formation is available.

The current study employed the sedimentation method adapted from Ledwoch et al. (2019) to form DSB on SS and PP coupons at 25 °C and the methodology of Harada and Nascimento (2021a, 2021b) to evaluate the sanitizer efficacy. The initial population (positive control) of *C. sakazakii* DSBs formed by both protocols ranged from 7.5 to 8.4 log CFU/cm² on surfaces tested with dry sanitizers. However, for coupons used for SH testing, there was an average difference of 1.7 log CFU/cm² in bacteria from DSB endpoint between PP and SS (8.0 and 6.3 log CFU/cm², respectively; data not shown). According to Nkemngong et al. (2020), a minimum DSB population density of 6.0 log CFU/coupon is required for sanitizer efficacy testing. All samples in the present study met or exceeded this threshold.

The five dry sanitizing methods showed similar performance (p < 0.05), with limited microbicidal action on both PP and SS surfaces, falling well below the performance criteria established by European and American standards, as mentioned above (UNE-EN 13697, 2024). Overall, no significant difference (p > 0.05) was observed among exposure times or between DSB protocols when evaluating for the same dry sanitizer. There was, however, a significant difference (p < 0.05) between surface materials against DSB from following T1, when treated with 70% ethanol for 5 min or hot air for 10 min. The desiccation stress experienced during DSB formation can alter stress-protein production, modify cell membrane composition, and promote the development of denser EPS matrices (Maillard & Centeleghe, 2023; Rahman et al., 2022). These factors likely contributed to the high resistance observed here against the dry sanitizing methods. In addition, according to Flemming & Wingender (2010), the EPS of biofilms subjected to desiccation stress has a greater number of non-specific binding sites, which can hinder sanitizer diffusion and reduce antimicrobial effectiveness.

Alcohol-based sanitizers are widely used in the LMF industry. For 70% ethanol, the greatest reduction in C. *sakazakii* viability (0.9 log CFU/cm²) was observed when DSB was formed with longer hydrated phases and shorter dry ones (T1) on PP after 5 min exposure, (Table 1). Meanwhile, the commercial product resulted in a maximum reduction in. bacteria in DSB of 0.6 log CFU/cm² after 15 min. There are currently no studies evaluating the effectiveness of alcohol-based sanitizers against *C. sakazakii* DSB. However, Jo et al. (2010) reported nearly 6-log CFU/coupon reductions in *C. sakazakii* hydrated biofilms using 70% ethanol. Lin et al. (2024) observed a reduction of 1.5 log CFU/sample of *Salmonella* DSB after 1 min exposure to isopropyl alcohol (70%). In contrast, Chaggar et al. (2022) noted a 5.9-log reduction in *S. aureus* dried hydrated biofilms after 1 min exposure to a commercial formulation containing 15% isopropanol + 7.5% ethanol + 0.76% QAC. In addition, the limited reductions achieved in the current study using alcohol-based sanitizers corroborate the previous report of Harsent et al. (2026) and Ledwoch et al. (2021), who highlight that the effective control of DSB requires the combination of chemical sanitizers with mechanical removal.

**Table 1.** Reductions in log CFU/cm² of dry surface biofilm (DSB) of *C. sakazakii* formed on polypropylene (PP) and stainless steel (SS) after sanitization for up to 30 min using: 70% ethanol (OH), a commercial product based on isopropyl alcohol (25%) and quaternary ammonium (0.015%) (CP), Hot air 90 ± 2 °C (HA), gaseous ozone 45 ± 2 mg/L (OZ), UV-C light 254 nm (UV), and sodium hypochlorite 200 mg/L, pH 6.5 (SH). Protocols were made with 2 cycles wet/dry phases: T1 (48/48) and T2 (24/120 h). Identical capital letters indicate that there was no statistical difference between averages in each column (exposure time) according to Tukey’s test (P<0.05). Identical lowercase indicates that there was no statistical difference between averages in each row (protocol and surface) according to Tukey’s test (P<0.05). Detection limit = 0.7 log CFU/cm².

| Protocol | Surface | Exposure time (min) [statistical comparison] |  |  |  |
| --- | --- | --- | --- | --- | --- |
|  |  | 5 | 10 | 15 | 30 |
| OH |  |  |  |  |  |
| T1 | PP | 0.9 ± 0.4 [Ba] | 0.3 ± 0.3 [Aa] | 0.6 ± 0.5 [Aa] | 0.5 ± 0.1 [Aa] |
|  | SS | 0.1 ± 0.4 [Aa] | -0.2 ± 0.2 [Aa] | -0.0 ± 0.4 [Aa] | 0.3 ± 0.1 [Aa] |
| T2 | PP | 0.3 ± 0.1 [ABa] | -0.0 ± 0.1.0 [Aa] | -0.0 ± 0.1 [Aa] | 0.1 ± 0.3 [Aa] |
|  | SS | 0.1 ± 0.2 [Aa] | 0.3 ± 0.3 [Aa] | 0.1 ± 0.2 [Aa] | 0.2 ± 0.2 [Aa] |
| CP |  |  |  |  |  |
| T1 | PP | 0.1 ± 0.1 [Aa] | 0.1 ± 0.2 [Aa] | 0.2 ± 0.3 [Aa] | 0.4 ± 0.3 [Aa] |
|  | SS | -0.0 ± 0.2 [Aa] | -0.0 ± 0.2 [Aa] | 0.6 ± 0.3 [Aa] | 0.5 ± 0.3 [Aa] |
| T2 | PP | -0.0 ± 0.2 [Aa] | -0.0 ± 0.1 [Aa] | 0.1 ± 0.4 [Aa] | 0.1 ± 0.2 [Aa] |
|  | SS | -0.0 ± 0.3 [Aa] | -0.0 ± 0.2 [Aa] | -0.1 ± 0.4 [Aa] | -0.1 ± 0.2 [Aa] |
| HA |  |  |  |  |  |
| T1 | PP | 0.3 ± 0.2 [Aa] | 0.6 ± 0.1 [Ba] | 0.5 ± 0.2 [Aa] | 0.5 ± 0.2 [Aa] |
|  | SS | 0.1 ± 0.4 [Aa] | -0.1 ± 0.1 [Aa] | 0.1 ± 0.3 [Aa] | 0.2 ± 0.5 [Aa] |
| T2 | PP | 0.1 ± 0.3 [Aa] | 0.1 ± 0.3 [ABa] | 0.1 ± 0.3 [Aa] | 0.3 ± 0.2 [Aa] |
|  | SS | -0.1 ± 0.1 [Aa] | -0.0 ± 0.2 [Aa] | 0.1 ± 0.3 [Aa] | -0.1 ± 0.1 [Aa] |
| OZ |  |  |  |  |  |
| T1 | PP | -0.0 ± 0.3 [Aa] | 0.1 ± 0.1 [Aa] | 0.1 ± 0.4 [Aa] | 0.2 ± 0.1 [ABa] |
|  | SS | 0.3 ± 0.3 [Aa] | 0.2 ± 0.2 [Aa] | -0.0 ± 0.4 [Aa] | -0.1 ± 0.4 [Aa] |
| T2 | PP | 0.2 ± 0.3 [Aa] | 0.4 ± 0.1.0 [Aa] | 0.1 ± 0.3 [Aa] | 0.6 ± 0.4 [Ba] |
|  | SS | 0.1 ± 0.3 [Aa] | 0.3 ± 0.5 [Aa] | 0.3 ± 0.5 [Aa] | 0.1 ± 0.1 [ABa] |
| UV |  |  |  |  |  |
| T1 | PP | 0.4 ± 0.9 [Aa] | 0.1 ± 0.3 [Aa] | 0.7 ± 0.5 [Aa] | 0.6 ± 0.7 [Aa] |
|  | SS | 0.1 ± 0.3 [Aa] | 0.2 ± 0.3 [Aa] | 0.2 ± 0.5 [Aa] | 0.3 ± 0.6 [Aa] |
| T2 | PP | 0.1 ± 0.5 [Aa] | 0.3 ± 0.4 [Aa] | 0.4 ± 0.1 [Aa] | 0.6 ± 0.1 [Aa] |
|  | SS | 0.2 ± 0.4 [Aa] | 0.3 ± 0.5 [Aa] | -0.0 ± 0.2 [Aa] | 0.3 ± 0.4 [Aa] |
| SH |  |  |  |  |  |
| T1 | PP | 1.3 ± 0.5 [Aa] | 2.6 ± 1.6 [ABa] | 3.4 ± 1.7 [Aa] | >7.2 ± 0.5 [Bb] |
|  | SS | 1.5 ± 1.3 [Aa] | 2.4 ± 0.7 [Aa] | 4.8 ± 0.4 [Ab] | >5.5 ± 0.1 [Ab] |
| T2 | PP | 1.5 ± 0.2 [Aa] | 2.0 ± 0.6 [Aa] | >7.3 ± 0.3 [Bb] | >7.3 ± 0.3 [Bb] |
|  | SS | 1.9 ± 0.6 [Aa] | 4.9 ± 0.1 [Bb] | 5.3 ± 0.9 [ABb] | 5.5 ± 0.7 [Ab] |

Hot air achieved reductions of ≤ 0.6 log CFU/cm², even after 30 min exposure (Table 1). The increased thermal resistance of DSB compared with wet surface biofilms (WSB) has been previously documented by Almatroudi et al. (2018). The authors reported reductions of 7.2 log CFU/coupon for WSB but only 0.9 log CFU/coupon for *S. aureus* ATCC 25923 DSB formed on polycarbonate after 10 minutes of exposure to 100 °C - values comparable to those observed in our study. These findings indicate that desiccation stress enhances the thermal tolerance of DSB. Bacterial thermal resistance is closely associated with the production of heat shock proteins (HSPs), particularly chaperone proteins, which help stabilize and refold damaged proteins under combined heat and low-moisture stress conditions. Recent studies have shown that pre-exposure to desiccation increases the expression of these chaperones and results in greater thermal tolerance in *C. sakazakii* (Wang et al., 2021; Zhu et al., 2024), although the expression of HSPs in DSB is yet to be documented.

Gaseous ozone and UV-C achieved maximum reductions of 0.6 and 0.7 log CFU/cm² after 30 and 15 min of exposure, respectively (Table 1). Epelle et al. (2024) studied UV light (185 nm, 3.22 mW/cm²), ozone gas (50 mg/L), and the combination of both against DSB of *C. auris* and *S. aureus* formed on polystyrene. After 20 min exposure, the authors noted significantly greater reductions than those obtained in the present study, > 4 and > 6 log CFU/mL on *S. aureus* and ca. 6 and 8 log CFU/mL on *C. auris*, for ozone and UV, respectively. This finding suggests that *C. sakazakii* DSB exhibits greater resistance to UV-C and ozone compared with other clinically relevant pathogens. In addition, Harada & Nascimento (2021a) evaluated the same sanitizers tested in the present study, on semi-hydrated biofilm of *Bacillus cereus* formed on PP and SS. The authors obtained better performance with UV-C and ozone when compared to our results, with reductions of 2 log CFU/cm² after 30 min. The superoxide dismutase enzyme has antioxidant capacity, mainly against superoxide radicals, and its action may be related to antimicrobial resistance to ozone (Sun et al., 2022). The gene encoding this enzyme has already been found in some *C. sakazakii* strains (Bao et al., 2017; Jin et al., 2024). The mechanisms underlying resistance to UV-C light have not yet been elucidated. However, Arroyo et al. (2012) suggest that high tolerance to UV-C could be associated with the production of yellow pigments by *C. sakazakii*, which would limit the action of the radiation on the bacterial cell. These mechanisms of resistance have yet to be investigated in *C. sakazakii* DSB.

The effect of SH was remarkably greater than that of dry treatments, with reductions above 1.0 log CFU/cm² after 5 min exposure. In addition, both exposure time and DSB protocol significantly influenced SH performance (p < 0.05), with reductions > 5.0 log CFU/cm² after 15 min when DSB were formed according to T2 and after 30 min for T1 (Table 1). These results suggest that the shorter wet-phase exposure (24 h) and/or the longer desiccation period (120 h) applied in the T2 protocol increased the sensitivity of *C. sakazakii* to SH. In contrast, Chaggar et al. (2022) reported greater resistance of *Pseudomonas aeruginosa* dried hydrated biofilms with 72-h dehydration compared to 24-h, reductions of 2.7 log CFU/coupon versus 3.8 log CFU/coupon after 1 min exposure to 0.39% SH. Interestingly, the initial action of SH on PP appeared to be slower than on SS; but over time, the reduction in microbial load on both surfaces reached the limit of detection (0.7 log CFU/coupon). Most available studies on DSB resistance to SH are focused on *S. aureus*. Ledwoch et al. (2019) reported a reduction of 5.8 log CFU/coupon after treatment with SH (1000 mg/L/2 min) in combination with mechanical action (10 s and 500 g pressure), using a wiperator. In contrast, other authors obtained a lower reduction in *S. aureus* DSB, ca. 3 log CFU/coupon, using the same SH concentration for 2-5 min exposure (Chowdhury et al., 2019; Ledwoch et al., 2021). Similar reduction was observed for *Salmonella* DSB after 1 min exposure to SH (Lin et al., 2024). In fact, our results indicate the need for longer exposure to SH to achieve reductions >4 log CFU/cm² in *C. sakazakii* DSB. This may be related to dense and compact DSB EPS (Rahman et al., 2022; Centeleghe et al., 2023), which may limit SH penetration into bacterial cells, thereby requiring longer contact times. This factor should be taken into account by the LMF industry when validating sanitation procedures. It is also important to highlight that, according to the recommendations of the Codex Alimentarius (CXC 66, 2008) the use of wet sanitization in PIF processing environments requires strict humidity control measures, with immediate drying of equipment and surrounding areas to ensure that any remaining water is removed immediately. Overall, further studies are needed to fully understand the mechanisms involved in DSB resistance to sanitizers, especially dry sanitizers, to enhance product performance in practice.

### 2.8. CLSM images

Figure 2 shows the CLSM images obtained for the DSB protocols after 0 and 30 min exposure to all sanitizers evaluated on both surfaces. Analysis of the images of the positive control samples (time 0) indicated DSB with heterogeneous coverage and percentage of green-stained cells (live), ranging from 37 to 75% (Figure 3). Lin et al. (2024) found a high percentage of red-stained cells (dead) in *Salmonella* DSB when compared to WSB, reaching 87% in DSB with 96 h. According to these authors, the outermost layers of DSB have a higher concentration of dead cells due to the greater desiccation stress to which they are exposed. In addition, these layers would serve as a protective barrier against antimicrobial agents for the innermost layers, which have a higher percentage of living cells. The same phenomenon of heterogeneity was reflected in the samples after 30 min exposure to sanitizers, which may have hindered the quantitative and qualitative analysis of CLSM images (Rumbaugh & Sauer, 2020). For 70% ethanol, there was a reduction in the percentage of green-stained cells in T1, but increase or no change was noted for T2 (Figure 3). On the other hand, the commercial product and ozone caused a reduction in the percentage of green-stained cells on both surfaces for DSB formed following either T1 or T2 protocols. For hot air, the same trend was observed for both DSB - on SS there was an increase in the percentage of green-stained cells and the opposite was observed on PP. Interestedly, in most scenarios evaluated for UV-C, there was an increase in the percentage of cells stained green after 30 min exposure (Figure 3). For SH, there was a difference between the DSB protocols. For T1, the percentage of green-stained cells remained stable on SS and increased on PP (from 58% to 75%), whereas for T2, it decreased on both surfaces—from 62% to 54% on SS and from 71% to 61% on PP (Figure 3). These observations align with the plate count results (Table 1), where T2 exhibited greater sensitivity to chlorine. However, since the counts were reduced to values close or below the detection limit, a high proportion of red-stained (dead) cells was expected in SH-treated samples. In fact, the high proportion of green-stained cells noted after SH treatment may suggest two feasible scenarios: overestimation of bacterial survival or presence of viable but non-culturable (VBNC) cells.

**Figure 2.**
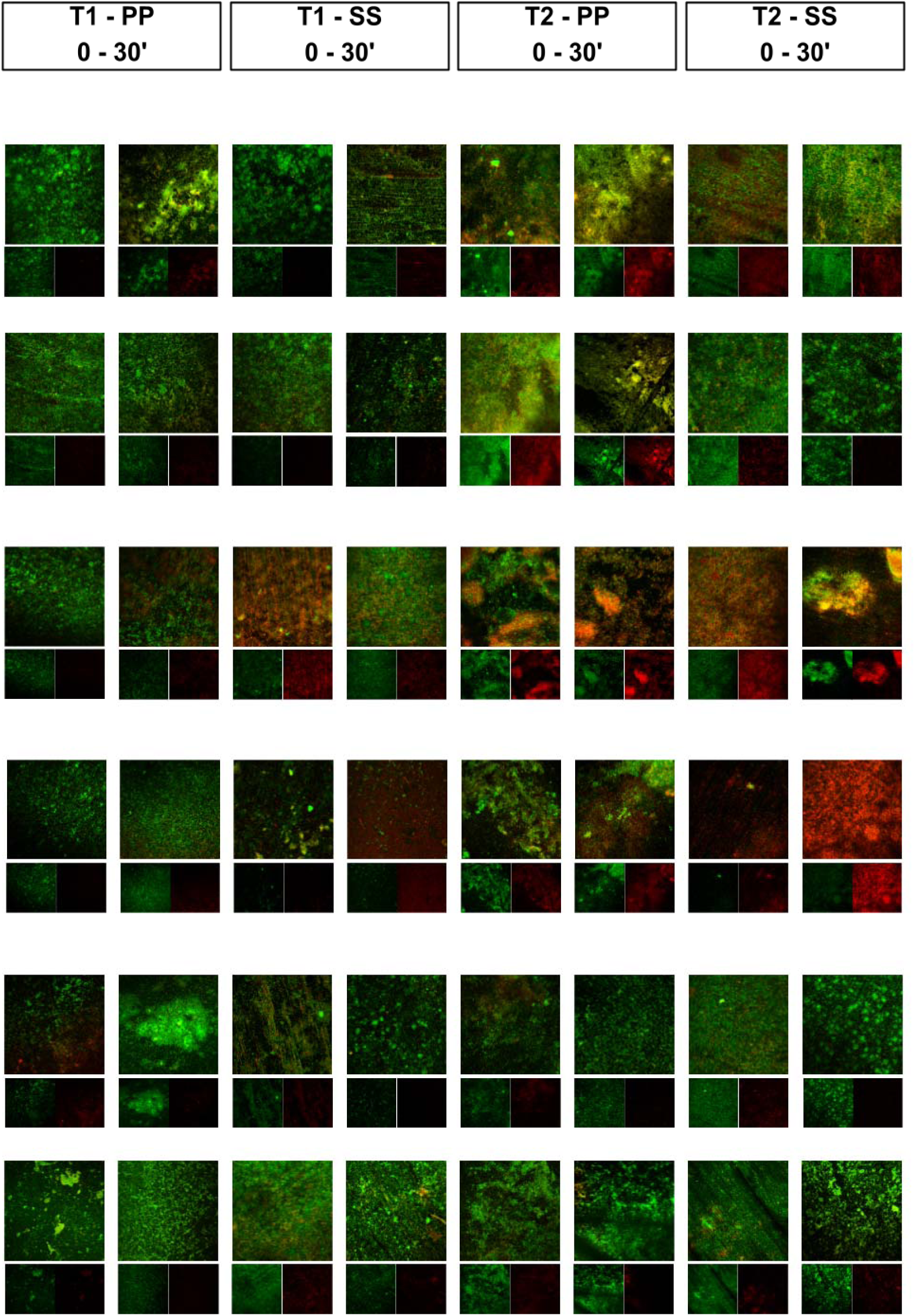
Confocal laser scanning microscopy (CLSM) images of *Cronobacter sakazakii* dry surface biofilm (DSB) formed on polypropylene (PP) and stainless steel (SS) before (0 min) and after 30-min sanitization. 70% ethanol (OH), a commercial product based on isopropyl alcohol (25%) and quaternary ammonium compound (0.015%) (CP), hot air (90 °C ± 2) (HA), gaseous ozone (OZ, 45 mg/L), UV-C light (254 nm) (UV), and sodium hypochlorite (SH, 200 mg/L pH 6.5). DSB protocols: T1 (48 h hydrated phase/48 h dry phase) and T2 (24 h hydrated phase /120 h dry phase). Green-stained cells (live) and red-stained cells (dead). Cells were stained using propidium iodide (dead) and SYTO® 9 (live), both at a dilution of 1:1000.

**Figure 3.**
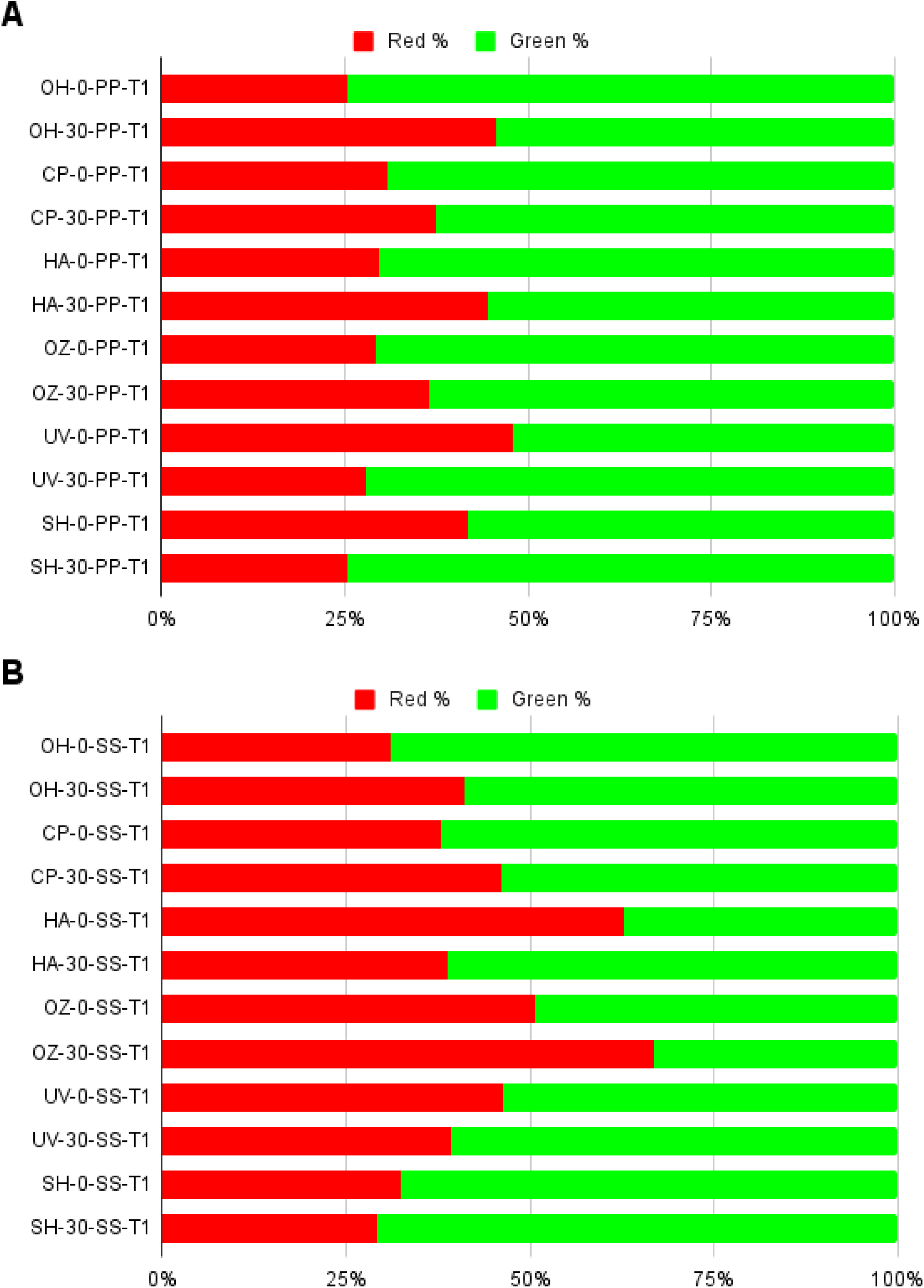
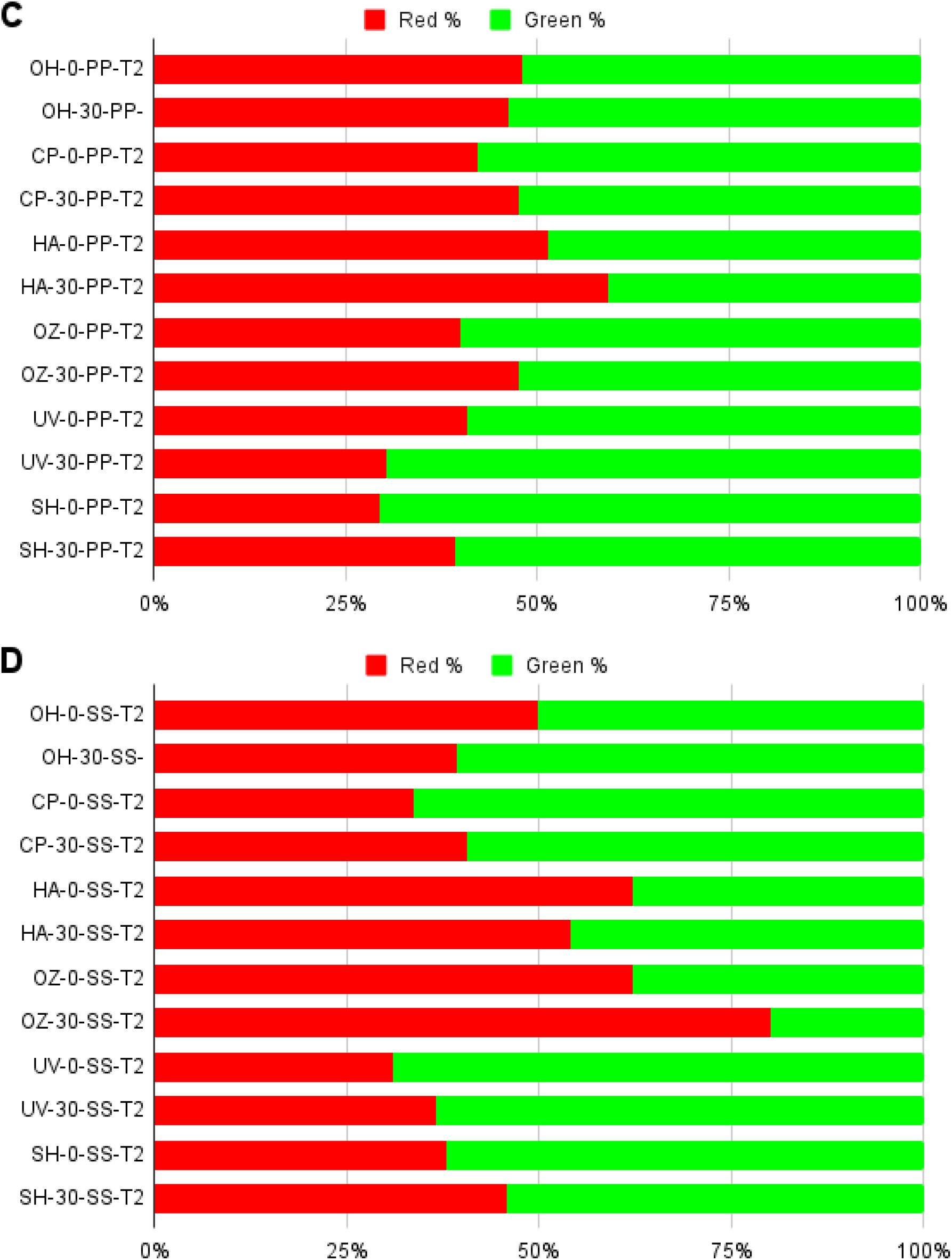
Percentage of green-stained cells (live) and red-stained cells (dead) obtained from CLSM using the LIVE/DEAD biofilm viability kit (composed of propidium iodide (dead) and SYTO® 9 (live), both at a dilution of 1:1000) of *Cronobacter sakazakii* dry surface biofilm (DSB) formed on polypropylene (PP) and stainless steel (SS) before (0 min) and after 30-min sanitization. 70% ethanol (OH), commercial product based on isopropyl alcohol (25%) and quaternary ammonium compound (0.015%) (CP), hot air (90 °C ± 2) (HA), gaseous ozone (OZ, 45 mg/L), UV-C light (254 nm) (UV), and sodium hypochlorite (SH, 200 mg/L pH 6.5). DSB protocols: T1 (48 h hydrated phase/48 h dry phase) and T2 (24 h hydrated phase /120 h dry phase). A: T1+PP; B: T1+SS; C: T2+PP; D: T2+SS.

The live/dead staining method is based on membrane integrity - Syto 9 permeates intact cells, whereas propidium iodide enters only cells with damaged membranes (Berney et al., 2007; Tchatchiashvili et al., 2025). For SH, inactivation of cellular metabolism (loss of culturability) occurs before significant membrane damage (Virto et al., 2005), meaning that, in this case, dead cells may fluoresce green, as observed in the current study. Luo & Raval (2025) reported that there is no direct correlation between the physiological state of cells indicated by live/dead dyes and cell morphology observed by scanning electron microscopy, suggesting an overestimation of bacterial survival when using dyes. Furthermore, according to Nocker et al. (2007), the concentration of propidium iodide required to obtain red-stained cells after exposure to chorine may be higher than usual. Therefore, complementary methods should be considered for a more accurate assessment of cell viability in DSB studies. Regarding VBNC cells, previous studies have demonstrated that *C. sakazakii* can enter a VBNC state in response to desiccation stress, including when in DSB (Jameelah et al, 2018; Yan et al., 2025; Vaz et al, 2026). Centeleghe et al. (2023) also reported higher viability than culturability in *Klebsiella* DSB aged 2 and 4 weeks, suggesting the presence of VBNC cells. As observed in the current study, Almatroudi et al. (2016) also noted a high proportion of green-stained cells after exposure of *S. aureus* DSB to 1,000–20,000 mg/L SH (pH 6.5–7.0) for 10 min, despite observing >7-log CFU reductions by plate count. Therefore, further studies based on molecular detection (PCR-PMA), flow cytometry, determination of metabolic activity or membrane integrity would be needed to confirm this hypothesis.

## 4. Conclusion

This study showed that DSB formed by *C. sakazakii* exhibited strong resistance to the dry sanitizing procedures, resulting in reductions of less than 1 log CFU/cm² over 30 min exposure. Indeed, from a risk management perspective, the control of *C. sakazakii* through dry sanitization is still a challenge for the LMF industry, especially for PIF processing plants. In contrast, SH achieved reductions greater than 5 log CFU/cm², highlighting its potential as an effective option for controlling *C. sakazakii* DSB in such environments, provided that strict humidity control and immediate drying of the production line after sanitization are ensured. However, CLSM images suggest the presence of cells in the VBNC state after chlorine sanitization, representing an additional challenge that must be considered when establishing environmental monitoring and control measures. Therefore, future studies should investigate the combined effects of different dry sanitization methods and their interaction with mechanical action. Additionally, studies focused on characterizing the metabolic and transcriptomic responses induced by desiccation stress, and how these responses contribute to increased tolerance to dry sanitizers, is essential to improve the understanding and control of these biofilms in industrial settings.

## 5. Acknowledgments

The authors thank the Fundação de Amparo à Pesquisa do Estado de São Paulo (Fapesp, process 2021/06809-2), and Coordenação de Aperfeiçoamento de Pessoal de Nível Superior – Brazil (CAPES; Finance Code 001). The authors also thank the National Institute of Science and Technology on Photonics Applied to Cell Biology (INFABIC) at the State University of Campinas (FAPESP, 2014/50938-8 and Conselho Nacional de Desenvolvimento Científico e Tecnológico, 465699/2014-6). Lastly, the authors thank the National Institute of Quality Control in Health, Oswaldo Cruz Foundation for donating the *C. sakazakii* strains.

## 6. Author contributions: CRediT

**Raul Fernando Pereira**: Investigation, Data curation, Formal analysis, Methodology, Writing – original draft; **Vinícius S. A. Vaz**: Investigation, Methodology; **Rafael Pimentel Maia**: Data curation, Formal analysis; **Jean-Yves Maillard**: Methodology, Writing – review and editing; **Maristela da Silva do Nascimento:** Conceptualization, Formal analysis, Funding acquisition, Methodology, Project administration, Resources, Supervision, Writing – review and editing.

## Confirmation of Publication and Licensing Rights

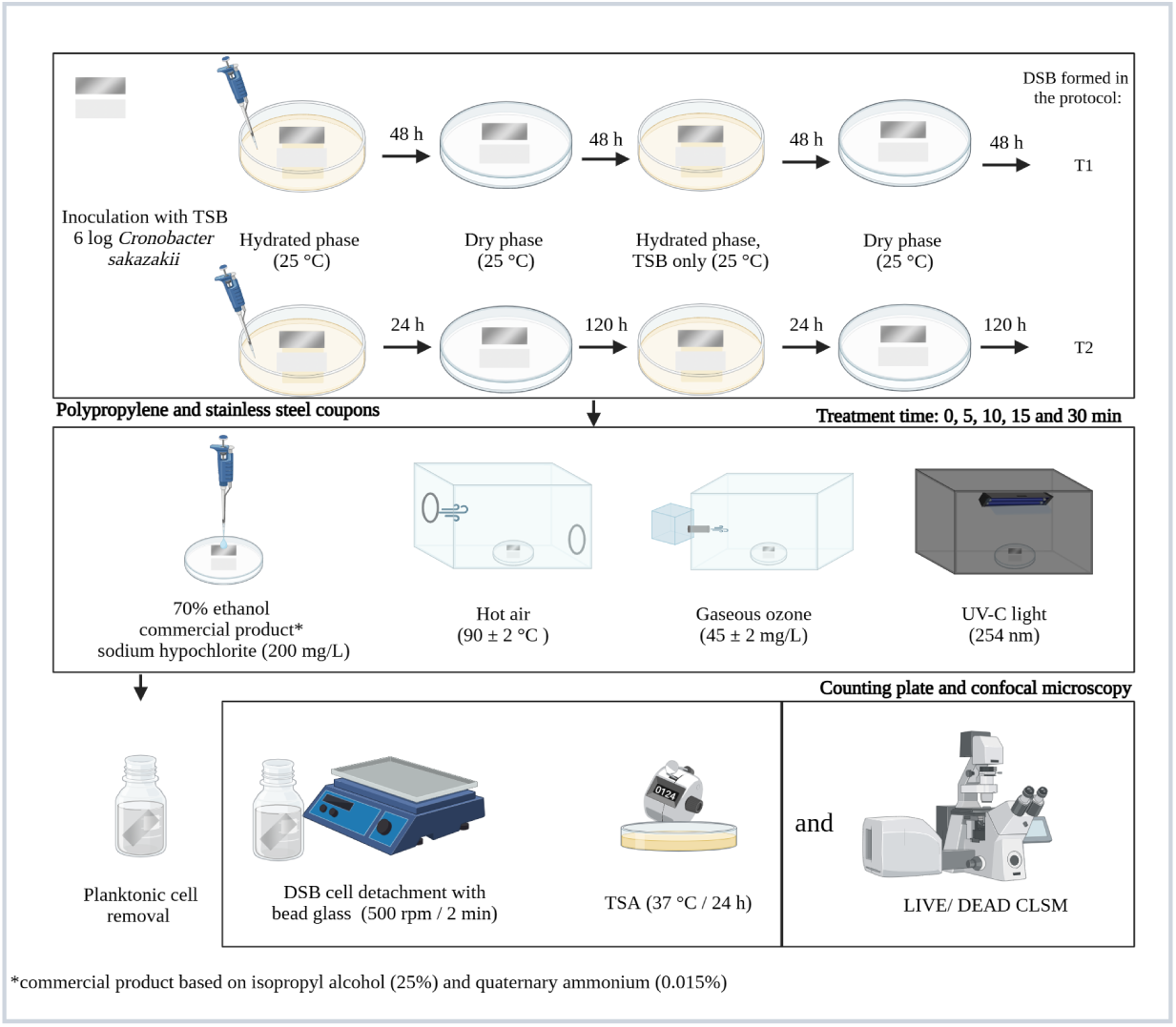

